# Age-related differences in electrophysiological correlates of visuospatial reorientation

**DOI:** 10.1101/2023.11.22.568209

**Authors:** Clément Naveilhan, Alexandre Delaux, Marion Durteste, Jerome Lebrun, Raphaël Zory, Angelo Arleo, Stephen Ramanoël

## Abstract

Spatial navigation abilities decline with age. Recent studies revealed a specific impairment in landmark-based reorientation, linked to changes in scene-selective brain regions activity. While fMRI studies suggest that these cortical modulations might be compensatory, a more precise investigation of the brain dynamics associated with visuospatial processing is warranted. We analyzed Event-Related Potentials and Event-Related Spectral Perturbations recorded from electrodes over scene-selective regions. 28 young adults and 28 older adults completed a desktop-based reorientation task using landmarks. Our findings show poorer reorientation performance among older adults. Signatures of age-related modulation of EEG activity imputable to scene-selective regions were predominantly observed within the right hemisphere. EEG analysis disclosed a tripartite worsening of scene processing accounting for older adults’ difficulties. Firstly, a delayed and reduced P1 component likely reflects a slower and less efficient stimulus discrimination. Secondly, an increased N1 amplitude and theta-band activity suggest a higher demand on cognitive resources associated with more effortful processing of visuospatial information. Thirdly, a decreased P2 amplitude may imply deficient attentional mechanisms to select task-relevant stimuli.

## 1. Introduction

Spatial navigation encompasses a complex set of behaviors that allow us to find our way and move in our environment. Although we are able to perform it effortlessly on a daily basis, successful navigation requires intricate cognitive processes such as sensory cue integration, working memory, or path integration (Wolbers & Hegarty, 2010), which are supported by a large and highly interconnected cerebral network (Ekstrom et al., 2017; Julian et al., 2018). Healthy aging is causally involved in the decline of spatial navigation abilities (Lester et al., 2017), with older adults experiencing difficulty in navigating both familiar and unfamiliar environments (Barrash, 1994). These impairments reduce the autonomy and mobility of older adults (Burns, 1999) resulting in an increased risk of progression of age-related disorders such as Alzheimer’s disease (Coughlan et al., 2018). Given the general aging of the population, it is essential to gain a better understanding of the factors contributing to these age-related navigational deficits and their neural correlates.

Among all the information required for spatial navigation, the ability to perceive and integrate visuospatial information plays a crucial role for humans who depend predominantly on their visual system to interact with their surroundings (Ekstrom, 2015; Foo et al., 2005). Vision allows humans to recognize the environmental context in which they are navigating and to rapidly encode the navigability of a visual scene presented (Greene & Oliva, 2009). The extraction of visual landmarks that provide information-rich cues for their orientation is also sustained by this sensory modality, allowing an efficient human spatial navigation (Fischer et al., 2020). Authors have observed that greater visual attention is indeed devoted to these landmarks, which subsequently serve as crucial reference points for successful navigational behavior (de Condappa & Wiener, 2014; Hamid et al., 2010; Wenczel et al., 2017). However, the ability to use landmark information for navigation declines with age, as evidenced by several studies (Harris & Wolbers, 2012; Hartmeyer et al., 2017; Wiener et al., 2012). More recently, Bécu et al. (2023) extended these findings by unveiling a specific decline in landmark-based navigation (*i.e.,* encoding objects) but a preserved performance during geometry-based navigation (*i.e.,* encoding spatial layouts). These two navigation modalities exhibit specific neural signatures in young adults (Ramanoël et al., 2022) highlighting the importance of considering their neural correlates to gain insight into the specific deficits of older adults in landmark navigation.

The integration of spatial visual cues is mediated by a network of high-level visual brain regions. This network comprises three scene-selective regions: the Parahippocampal Place Area (PPA) (Epstein & Kanwisher, 1998), the Retrosplenial Complex (RSC, also referred as Medial Place Area, MPA) (Maguire, 2001; Silson et al., 2016), and the Occipital Place Area (OPA) (Dilks et al., 2013). The PPA, located in the parahippocampal cortex, is thought to be involved in the representation of the spatial layout (Kravitz et al., 2011) and in landmark recognition (Janzen & van Turennout, 2004; Sun et al., 2021), thus contributing to scene categorization (Persichetti & Dilks, 2018). The RSC, a region of the posterior cingulate cortex, is involved in the computation of heading directions (Gramann et al., 2021; Marchette et al., 2014), the translation of information between egocentric and allocentric spatial reference frames (Vann et al., 2009; Zhang et al., 2012), and it may also combine visual and motor inputs for landmark encoding (Fischer et al., 2020). Finally, the OPA, which is located near the transverse occipital sulcus, supports first-person vision-guided navigation through its role in encoding environmental boundaries, local elements, and potential paths in a scene, which are also called navigational affordances (Bonner & Epstein, 2017; Epstein et al., 2017; Julian et al., 2018).

In the context of aging, several functional magnetic resonance imaging (fMRI) studies have highlighted age-related modifications in scene-selective regions during visual and spatial processing. Notably, reduced activity in the PPA has been observed to underpin age-related differences in the categorization of the fine-grained content of visual scenes (Ramanoël et al., 2015). Furthermore, the neural specificity and distinctiveness of the PPA and RSC have been shown to decline with age and to predict individual source and spatial memory abilities (Koch et al., 2020; Srokova et al., 2020). Regarding the OPA, fMRI acquisitions during a Y-maze reorientation task using objects as landmarks showed an increased activity of this region in older adults (Ramanoël et al., 2020). Critically, this age-related increase in parietal activity was only reported during the reorientation phase of the task involving landmark processing but not during free navigation (not involving reorientation). This finding was complemented by another brain connectivity study reporting a preserved structural connectivity around the OPA region and an increased functional connectivity between the OPA and PPA in older adults (Ramanoël et al., 2019).

Despite these results, how the temporal dynamics of scene-selective regions could contribute to the age-related navigational decline remains poorly characterized, mainly due to the limitations of fMRI in capturing brain processes at the millisecond timescale (Glover, 2011). With its high temporal resolution, Event-Related Potential (ERP) analysis represents a valuable neuroimaging approach for investigating early perceptual processes with electroencephalography (EEG). Notably, one ERP component, the occipito-parietal P2, has been proposed to be a marker of scene processing (Harel et al., 2016) and to reflect the activity of scene-selective regions (Kaiser et al., 2020). These results were complemented with intracranial EEG recordings suggesting that the activity of the OPA occurs in the time period of the P2 component (Vlcek et al., 2020). Recently, the amplitude of the P2 component was reported to scale linearly with the number of navigational affordances (Harel et al., 2022) reinforcing the P2 as a marker of scene-selective activity and more specifically of the OPA. Based on recordings from these occipito-parietal electrodes, Lithfous et al. (2014) reported an age-related increase of the P2 component amplitude and delayed P2 latency associated with an impaired performance on a spatial localization task. They suggested that changes of the parietal P2 component may reflect the mechanisms underlying the age-related decline in spatial processing and they emphasized the need for further studies to investigate P2 in relation to spatial memory or spatial visual cue processing. In a subsequent study using an EEG time-frequency analysis, Lithfous et al. (2018) found an increase in parahippocampal *theta* activity in high-performing older adults compared to young adults during a spatial navigation task. They also found a decreased *theta* power in the group of low-performing older adult (*i.e.,* the group with the lowest accuracy in reproducing paths), reinforcing the proposed relationship between parahippocampal *theta* oscillations and successful navigation (Bohbot et al., 2017; Chrastil et al., 2022; Jacobs, 2014). These results highlight the potential of using EEG (Event Related Potential and time frequency analyses) to investigate the neural dynamics associated with reorientation impairment in older adults. However, none of these studies considered age-related differences in visuospatial processing despite the considerable impact of age on this cognitive function (Bécu et al., 2023; Segen et al., 2021). Indeed, aging is associated with declines in visual acuity (Faubert, 2002) and a reduced capacity for fine processing which may partially account for navigational impairments in older adults, even more so in environments where visual landmarks are the sole cues for reorientation (Ramanoël et al., 2015, 2020).

To address this caveat, the present study aims to examine age-related differences during a landmark-based reorientation task and the associated brain dynamics using EEG recordings from electrodes related to scene-selective brain regions (Harel et al., 2016; Kaiser et al., 2020). In order to investigate the contribution of age-related visuospatial processing declines in reorientation, we manipulated the level of perceptual difficulty, leading to the presentation of large and small landmarks. We hypothesized that older participants would exhibit a poorer navigational performance than young participants, especially when perceptual difficulty is increased (*i.e.,* when landmarks are smaller). At the cortical level, we expected that older adults would show higher parietal P2 amplitude and *theta* activity during reorientation than young adults, reflecting an increased involvement of the scene-selective regions.

## 2. Methods

### 2.1 Participants

We conducted the experiment on a sample population of 30 young participants and 32 healthy older participants. We removed 2 older participants from the analysis because they performed below chance level, and then we cannot ensure their comprehension of the task. In addition, 4 other participants (2 older and 2 young participants) were excluded due to excessive artefacts in the EEG data as assessed by signal-to-noise ratio calculation and a careful visual inspection of the signals. Analyses were finally conducted on 28 young participants (mean age: 23.93 years old; SEM = 0.64; range: 19-35; 14 females) and 28 older adults (mean age: 71.25 years old; SEM = 1.01; range: 61–81; 18 females). Participants were right-handed, had no history of neurological or psychiatric disorders and they self-reported normal or corrected-to-normal vision. They were assessed for cognitive impairment using the GRECO French version of the MMSE (Kalafat et al., 2003) with the proposed 26 cut-off to ensure their healthy cognitive status. They also completed a computerized version of the Spatial Orientation Task (Friedman et al., 2020). These results are presented and discussed in the Supplementary Materials (**Table S1**). The experiment was approved by the local Ethical Committee (CERNI-UCA no. 2021-050) and participants provided informed consent before starting the experiment.

### 2.2 Stimuli and procedure

Visual stimuli were created using the Unity Engine software (Unity Technologies version 2019.2.0.0f1) and presented on an iiyama ProLiteB2791HSU monitor (1920×1080, 30-83khz) placed at eye level and 60 cm away from the participants. Stimuli were presented using the open-source PsychoPy software (v2022.13), implemented on a Dell Precision 7560 computer (Intel® Xeon® W-11955).

The environment was adapted from a previous fMRI experiment on healthy aging (Ramanoël et al., 2020). It was a three-arm maze (Y-maze) consisting of three corridors: one branch containing a goal materialized by a gift box, 2 identical starting branches, and 3 three-dimensional (3D) objects positioned at the intersection serving as landmarks (a cube, a ball, and a pyramid). The experimental paradigm was divided into 3 tasks: learning, reorientation, and control (**Figure 1**). During the learning task, participants were passively moved through the maze at 2.5 virtual meter per second, with a rotation speed of 40 rad/s. They were instructed to memorize the path to the goal using the objects positioned at the intersection. Then, during the reorientation task, participants were presented with images of the intersection taken from the videos, and they were instructed to indicate the direction of the goal, as quickly and accurately as possible using their right hand to press the directional keys (left or right). These snapshots were extracted from either a near (at 4.25 virtual meters from the intersection) or a far perspective (at 11.2 virtual meters from the intersection) to modulate perceptual difficulty (hereafter referred to as *large* and *small* conditions, respectively). The average retinal visual angle of the landmarks in the small condition was 1.2°, while it was 2.5° in the large condition. Images were presented in a pseudo-randomized order (*i.e.,* a similar stimulus was presented no more than three times in a row) for 3 seconds each and were followed by auditory feedback depending on the correctness of the answer given. Afterwards, participants performed the control task which consisted of passively viewing images of the intersection, but this time with all 3 objects being identical (3 spheres, 3 cubes or 3 pyramids). They were instructed to look carefully at both the objects and the fixation cross presented between the different images. To mitigate the possibility of participants losing interest, we varied some properties of the environment (*i.e.,* wall texture and goal location), leading to the presentation of 15 different combinations, presented pseudo-randomly across participants. This sequence of learning, reorientation and control tasks was repeated 3 times for a total of 60 videos, 300 reorientation trials and 180 control trials, and a total acquisition time of 49 minutes.

**Figure 1.**
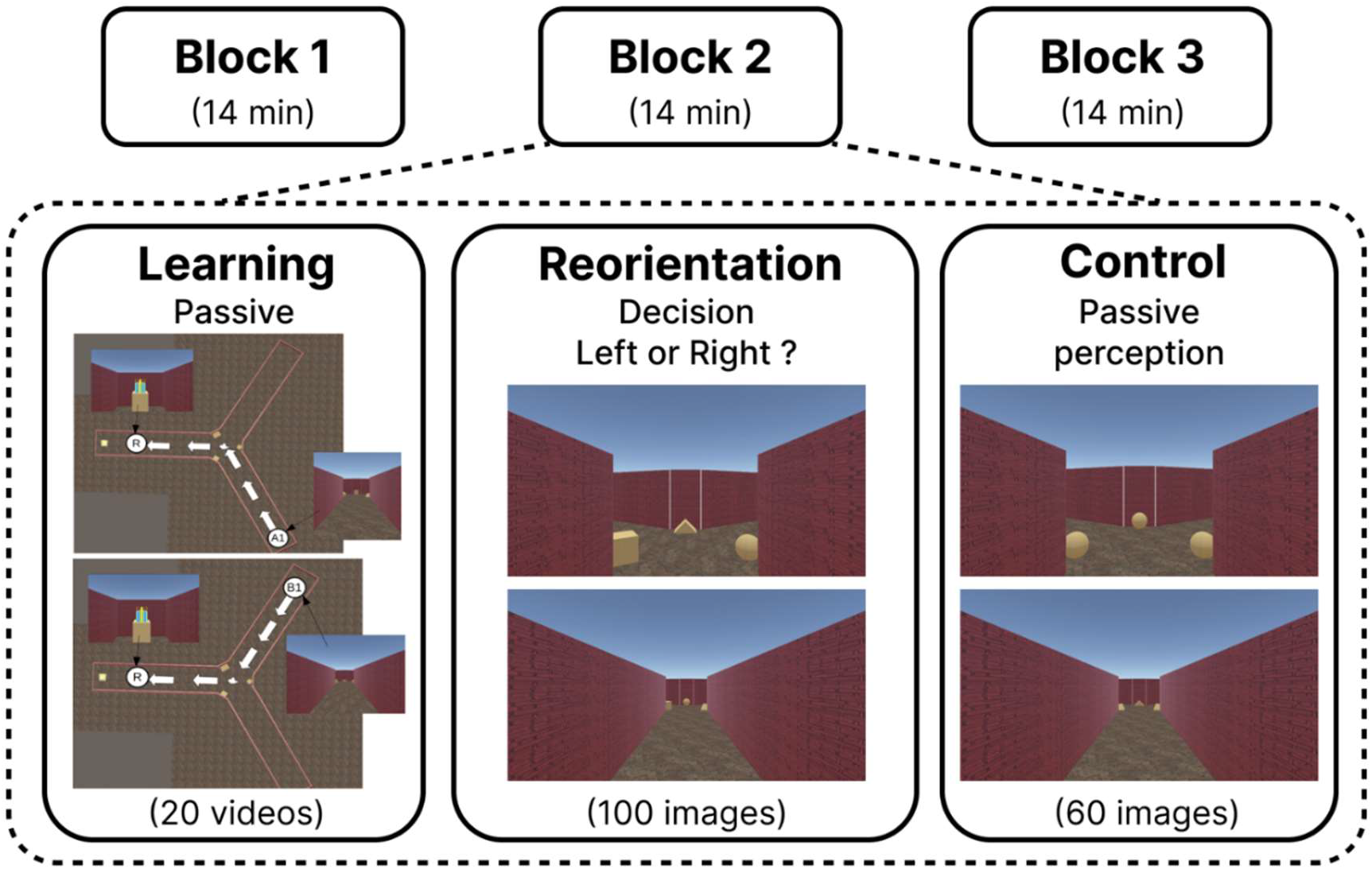
Presentation of the paradigm and some of the stimuli used. Blocks were the same between participants but were presented in a randomized order, with different wall textures. A short break was proposed to all participants between blocks

### 2.3 Recording and Analysis

#### 2.3.1 EEG recording and preprocessing

EEG was sampled at 500 Hz from 64 Ag/AgCl electrodes mounted in a cap (Waveguard™ original) connected to an amplifier (eego™ mylab, ANT Neuro) and digitized at 24-bit resolution. Data were referenced to CPz electrode, with AFz as ground. Electrode impedances were reduced to at least 20 kΩ, with most values falling below 10 kΩ. Synchronization of EEG recording, and stimulus presentation was ensured by the LabStremingLayer software Labrecorder (LSL Labrecorder version 1.13). Offline preprocessing was performed with MATLAB (R2021a; The MathWorks Inc., Natick, MA, USA) using custom scripts based on the EEGLAB toolbox version 14.1.0b (Delorme & Makeig, 2004) and adapted from a previously used processing pipeline (Delaux et al., 2021).

We first downsampled the data to 250 Hz and corrected the time points for the software delay using the trigger time added to a fixed delay of 50ms for the hardware delay. We automatically removed line noise using the recently developed Zapline-Plus algorithm (Klug & Kloosterman, 2022). We then automatically identified and rejected noisy channels, using the default parameters proposed in the PREP pipeline (Bigdely-Shamlo et al., 2015). On average, 4.25 channels were rejected (SEM= 0.39). These channels were then reconstructed by spherical interpolation of neighboring channels, and the data were re-referenced to the common average. Artifacts were automatically rejected using Artifact Subspace Reconstruction (ASR) (Kothe & Jung, 2015) which uses clean portions of the data to determine thresholds for rejecting components. We used an ASR cutoff parameter of 20, corresponding to the proposed optimal range between 10 and 100 (Chang et al., 2020). We then temporally high-passed the cleaned dataset using a 1.5 Hz Finite Impulse Response (FIR) filter with a Hamming window (with 0.5 Hz transition bandwidth, 1.25 Hz passband edge, and 1650 order) (Klug & Gramann, 2021) before applying independent component analysis (ICA) using the Adaptative Mixture Independent Component Analysis (AMICA) algorithm (Palmer et al., 2008). Next, each independent component (IC) was assigned a dipolar source reconstructed with the equivalent dipole model (DipFit; Oostenveld & Oostendorp, 2002). We used the ICLabel algorithm (Pion-Tonachini et al., 2019) to classify components into 7 classes, using default percentages for classification. We opted for a conservative approach and on average, we retained 14.64 components (SEM = 0.67) corresponding to brain activity. Next, we applied a bandpass filter to the data, with a lower cutoff frequency of 0.3 Hz (with 0.5 Hz transition bandwidth, 0.55 Hz passband edge, and 1650 order) to remove slow drifts, and an upper cutoff frequency of 80 Hz (20 Hz transition bandwidth, 80 Hz passband edge, and 42 order) to attenuate high-frequency noise and muscle artifacts. Finally, the preprocessed data were segmented into epochs ranging from -200 ms before to 600 ms after stimulus onset for all conditions. Epochs were excluded from the analysis if less than 90% of the period was clean. On average, we kept 76.56% of the epochs (mean epochs kept per subject: 367.50; SEM = 2.60). The number of epochs extracted was the same for both age groups (t_(1,43.33)_ = 0.820, *p* = 0.417).

#### 2.3.2 Event Related Potential and time-frequency analyses

Analyses were restricted to the occipito-parietal electrodes previously associated with scene-selective regions (Harel et al., 2016; Kaiser et al., 2020), corresponding to P6-P8-PO8 for the right hemisphere and P5-P7-PO7 for the left hemisphere. Further data analysis was performed using custom MATLAB scripts with functions from the Fieldtrip toolbox (Oostenveld et al., 2011).

For the Event Related Potentials (ERPs), the baseline was identified from -200 ms to image onset corresponding to the recommended minimum of 10 to 20% total epoch length (Luck, 2014), and mean baseline activity was subtracted from the data. Peak amplitude was calculated using the mean amplitude, corresponding to the average of the most positive value surrounded by two lower values. Given the reported effect of aging on peak latency (Kropotov et al., 2016; Mueller et al., 2008), we decided to calculate P1, N1, and P2 latency for each age group. Then, we took a 100 ms wide window around this value to extract the latencies and amplitudes of the individual components.

Time-frequency analysis was performed using the superlet approach (Moca et al., 2021), a spectral estimator that uses sets of increasingly constrained bandwidth wavelets to achieve time-frequency super-resolution. For this purpose, we used the Fieldtrip function ft_freqanalysis to decompose between 1 and 80 Hz, using a width of 2 and a Gaussian width of 3, with an increasing order scaling from 1 to 80. Once the decomposition was completed, we computed event-related spectral perturbations (ERSPs) (Makeig, 1993). We used the decibel conversion to normalize power values with a baseline between -250 and -50 ms due to temporal smearing (see Cohen, 2014 for more details).

#### 2.3.3 Statistical analysis

For the ERSPs, we performed a non-parametric cluster-based permutation test using a Monte-Carlo estimate to deal with the multiple comparisons problem (Maris & Oostenveld, 2007). We chose the most robust and least conservative method among several modalities, which involved 10 000 permutations with weighted cluster mass (Hayasaka & Nichols, 2004) and a cluster-level *alpha* of 0.005 to account for multiple comparisons. All other statistical analyses were performed using R Statistical Software (version 4.2.1, R Foundation for Statistical Computing, Vienna, Austria) with R studio (version 2022.07.02+576). After comparing different models using the Akaike information criterion (Akaike, 1974), we decided to use the linear mixed model from the lme4 R package (Bates et al., 2014), with participants included as random intercept. Then, we used the anova function to compute a type III Analysis of Variance (ANOVA) using Satterthwaite’s method. The reported results are estimated marginal means computed with the emmeans package in R, using a type III sum of squares, and finally post-hoc Tukey’s Honest Significant Difference tests were performed. To ensure normality of residuals and homoscedasticity, both were carefully visually inspected using quantile-quantile plots and boxplots, respectively. Finally, we conducted correlation analyses between ERP peaks and behavioral data using the Bonferroni correction to account for multiple comparisons.

## 3. Results

### 3.1 Behavioral results

We observed age-related differences in navigation performance. In terms of accuracy **(Figure 2.A)**, we reported only a main effect of age (F_(1,54)_ = 6.63, *p* = 0.013, η_p_^2^ = 0.11, 95% CI = [0.00, 0.28])), with lower accuracy for older adults (M = 93.4%, SE = 0.94) compared to young adults (M = 96.8%, SE = 0.94). There was neither an effect of condition (F_(1,54)_ = 0.527, *p* > 0.47) nor an interaction between the factors (F_(1,54)_ = 0.136, *p* > 0.71). Regarding the reaction time (**Figure 2.B**), we found a main effect of age (F_(1,54)_ = 40.97, *p* < 0.001, η_p_^2^ = 0.43, 95% CI = [0.24, 0.58]) in older adults (M = 1162 ms, SE = 35.2) who had a longer reaction time than young adults (M = 843 ms, SE = 35.2). We also found a main effect of condition (F_(1,54)_ = 52.47, *p* < 0.001, η_p_^2^ = 0.48, 95% CI = [0.30, 0.63]) with a higher reaction time for the small condition (M = 1020 ms, SE = 25) compared to the large condition (M = 985 ms, SE = 25) for both young and older adults. There was no interaction between age and condition (F_(1,54)_ = 1.48, *p* = 0.229).

**Figure 2.**
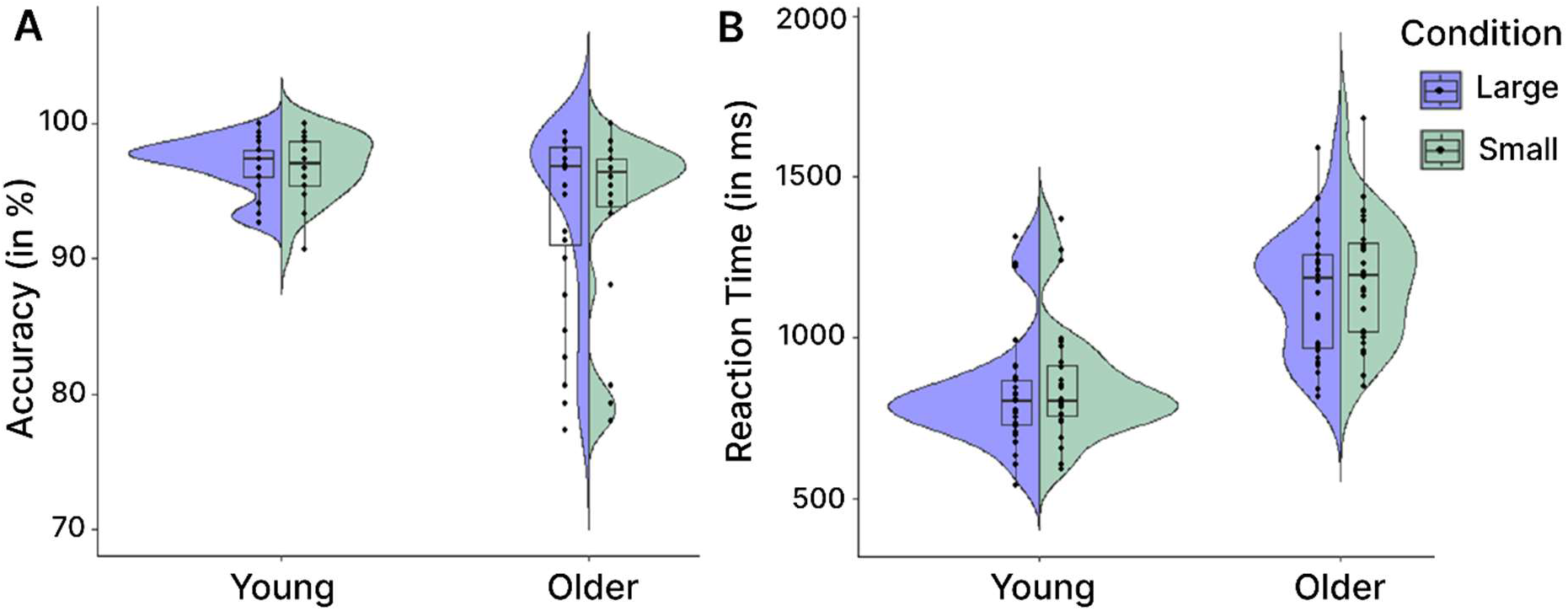
Performance of participants in the reorientation task. Individual points represent the average for each participant. **A**. Accuracy computed as the percentage of reorientation errors over all trials. **B.** Reaction time between the presentation of the stimulus and the recorded response.

### 3.2 ERPs results for large and small conditions during reorientation

In this analysis we compared the large and small conditions to examine the effect of age with perceptual difficulty during the reorientation task (**Figure 3**).

**Figure 3.**
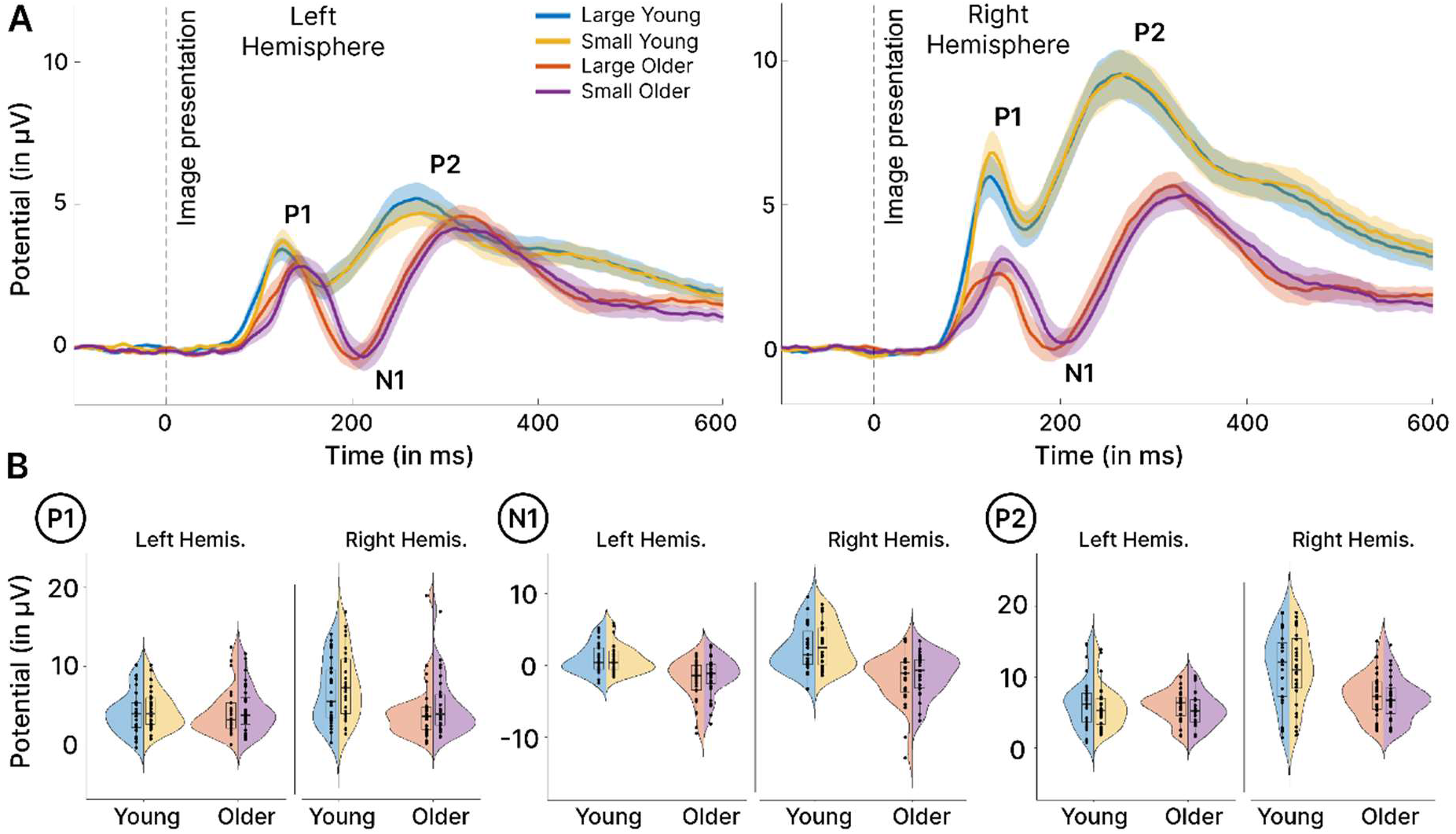
ERP results for near and far conditions during the reorientation task. **A.** Grand-average ERPs considering age (young/older), condition (large/small) and hemisphere (left/right) as variables, individually baselined corrected. Activity averaged over P6-P8-PO8 electrodes for the right hemisphere and over P5-P7-PO7 electrodes for the left hemisphere. **B.** Split violin plot of extracted individual amplitudes for P1, N1 and P2 component. Statistics were computed using these values.

#### P1, N1, and P2 amplitudes

First, we evaluated the age effect by comparing the EEG recordings of young and older adults. We found no main effect of age for P1 amplitude (F_(1,54.17)_ = 2.57, *p* = 0.11), but a higher amplitude for young than older adults when considering only the right hemisphere (t_(67)_ = 3.02, *p* = 0.018, d = 1.29, 95% CI = [0.43, 2.15]). We reported an increased N1 amplitude (*i.e.,* more negative) for older adults compared to young adults (F_(1,54.09)_ = 28.92, *p* < 0.001, η_p_^2^ = 0.35, 95% CI = [0.16, 0.51]) in both hemispheres. We also observed a lower P2 amplitude in older adults compared to young adults (F_(1,54)_ = 7.36, *p* < 0.001, η_p_^2^ = 0.12, 95% CI = [0.01, 0.29]) but this age effect was restricted to the right hemisphere (t_(68)_ = 4.58, *p* < 0.001, d = 1.82, 95% CI = [-1.00, 2.62]), with no difference for the left hemisphere (t_(68.3)_ = 0.53, *p* = 0.95).

Next, we considered the effect of condition comparing large *vs.* small. We found a higher P1 amplitude for the small condition compared to the large condition (F_(1,159.25)_ = 4.91, *p* = 0.028, η_p_^2^ = 0.03, 95% CI = [0.00, 0.10]). There was no modulation of either N1 (F_(1,159)_ = 3.43, *p* = 0.066), or P2 amplitudes (F_(1,162)_ = 1.70, *p* = 0.19).

Finally, we examined the lateralization effect by directly comparing the brain signals from the right and left hemispheres. We reported a higher amplitude for P1 in the right hemisphere (F_(1,159.77)_ = 33.74, *p* < 0.001, η_p_^2^ = 0.17, 95% CI = [0.08, 0.28]), but this effect was restricted to young adults, as older adults showed no lateralization (t_(159)_ = 0.76, *p* = 0.87). For P2, we found a higher amplitude in the right hemisphere independent of age group (F_(1,162)_ = 122.29, *p* < 0.001, η_p_^2^ = 0.43, 95% CI = [0.32, 0.52]). Considering N1, we observed the opposite, with a greater amplitude in the left hemisphere (F_(1,159.79)_ = 15.15, *p* < 0.001, η_p_^2^ = 0.09, 95% CI = [0.02, 0.18]). This effect was restricted to young adults (t = 5.13, *p* < 0.001, d = 0.99, 95% CI = [0.60, 1.39]), whereas older adults presented similar N1 activity for both hemispheres (t_(159)_ = 0.32, *p* = 0.99).

#### P1, N1, and P2 latencies

Looking at the age effect, we found later peaks for older adults in P1 (F_(1,54.35)_ = 13.12, p < 0.001, η_p_^2^ = 0.19, 95 % CI = [0.04, 0.37]), N1 (F_(1,53.92)_ = 37.24, *p* < 0.001, η_p_^2^ = 0.41, 95 % CI = [0.21, 0.56]) andP2 (F_(1,54)_ = 130.25, *p* < 0.001, η_p_^2^ = 0.71, 95% CI = [0.57, 0.79]). The effect for P1 was restricted to the small condition (t_(77)_ = 4.76, *p* < 0.001, d = 1.57, 95% CI = [0.90, 2.24]) as we reported no age difference for the large condition (t_(77)_ = 1.85, *p* = 0.26).

Considering the condition effect, we observed a delayed P1 for the small condition compared to the large condition (F_(1,159.53)_ = 20.89, *p* < 0.001, η_p_^2^ = 0.12, 95% CI = [0.04, 0.21]). This effect was not present for either N1 (F_(1,159.28)_ = 1.85, *p* = 0.544), or P2 (F_(1,162)_ = 1.52, *p* = 0.22). This condition effect for P1 was present only for older adults (t_(159)_ = 5.80, *p* < 0.001, d = 1.10, 95% CI = [0.71, 1.49]) with no difference for young adults (t_(159)_ = 0.70, *p* = 0.90).

Finally, concerning lateralization, we found no effect for P1 (F_(1,160.32)_ = 3.76, *p* = 0.054) or N1 (F_(1,160.41)_ = 3.64, *p* = 0.058). For P2, we observed a later peak in the right hemisphere compared to the left hemisphere (F_(1,162)_ = 4.14, *p* = 0.043, η_p_^2^ = 0.03, 95% CI = [0.00, 0.54]).

### 3.3 ERP results comparing reorientation and control tasks

In this second analysis, we compared the activity elicited by the reorientation task and the control task, in order to dissociate reorientation from pure scene perception effects. The results presented below correspond to the large and small conditions merged together (**Figure 4**). Before doing so, we checked that the same pattern of results was obtained when the two conditions were considered separately.

**Figure 4.**
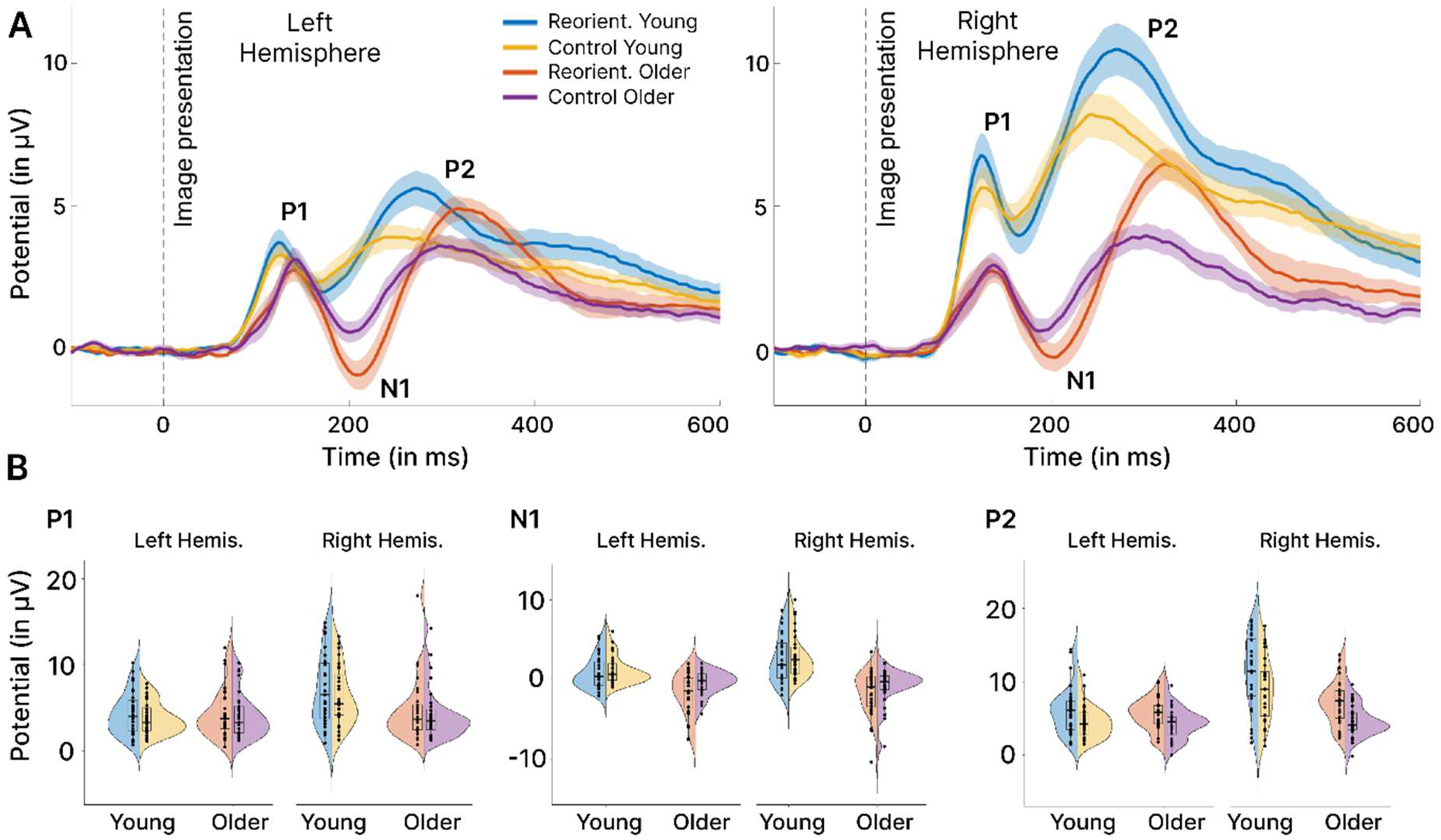
ERP results comparing reorientation and control tasks. **A.** Grand-average ERPs considering age (young/older), task (reorientation/control) and hemisphere (left/right) as variables, individually baselined corrected. Activity averaged over P6-P8-PO8 electrodes for the right hemisphere and over P5- P7-PO7 electrodes for the left hemisphere. **B.** Split violin plot of extracted individual amplitudes for P1, N1 and P2 components. Statistics were computed using these values.

#### P1, N1 and P2 Amplitudes

First, we examined the age effect, comparing young and older adults. We found no effect for P1 amplitude (F_(1,54.06)_ = 2.15, *p* = 0.15), but a higher amplitude for young adults when considering the right hemisphere only (t_(68)_ = 2.99, *p* = 0.019, d = 1.21, 95% CI = [0.39, 2.02]), with no age difference in the left hemisphere (t_(67.6)_ = 0.23, *p* = 0.996). We found a higher N1 amplitude (*i.e.,* more negative) for older adults (F_(1,54.08)_ = 31.82, *p* < 0.001, η_p_^2^ = 0.37, 95% CI = [0.18, 0.53]), and the opposite pattern for P2, with a higher amplitude for young adults (F_(1,84)_ = 10.33, *p* = 0.002, η_p_^2^ = 0.16, 95% CI = [0.02, 0.34]).

We then considered the task and compared reorientation and control (*i.e.,* passive perception). We found a similar pattern, with a higher amplitude for the reorientation task for P1 (F_(1,161.14)_ = 4.32, *p* = 0.040, η_p_^2^ = 0.03, 95% CI = [0.00, 0.09]), N1 (F = 13.90, *p* < 0.001, η_p_^2^ = 0.08, 95% CI = [0.02, 0.17]) and P2 (F_(1,162)_ = 44.38, p < 0.001, η_p_^2^ = 0.22, 95% CI = [0.11, 0.32]). Considering only N1 amplitude, the effect was limited to older adults (t_(159)_ = 3.98, *p* < 0.001, d = 0.75, 95% CI = [0.37, 1.14]), as young adults showed no difference between reorientation and control for this component (t_(161)_ = 1.30, *p* = 0.57).

Finally, we considered laterization, comparing the left and right hemispheres. We reported a similar pattern for positive components, with a higher amplitude in the right hemisphere for P1 (F_(1,161.14)_ = 32.22, *p* < 0.001, η_p_^2^ = 0.17, 95% CI = [0.07, 0.27]) and P2 (F = 95.95, *p* < 0.001, η_p_^2^ = 0.37, 95% CI = [0.26, 0.47]). These results were only observed for young adults, as older adults showed no lateralization effect for either the P1 (t_(161)_ = 0.58, *p* = 0.94) or the P2 component (t_(162)_ = 2.43, *p* = 0.12). For N1, we found the opposite result, with a higher amplitude for the left hemisphere (F_(1,161.16)_ = 14.83, *p* < 0.001, η_p_^2^ = 0.08, 95% CI = [0.02, 0.18]), again only for young adults (t = 5.94, *p* < 0.001, d = 1.13, 95% CI = [0.73, 1.52]) with no lateralization for older adults (t_(161)_ = 0.51, *p* = 0.96).

#### P1, N1 and P2 Latencies

First, we observed a similar pattern for age differences across components, with a later peak for P1 (F_(1,54.06)_ = 12.12, *p* < 0.001, η_p_^2^ = 0.18, 95% CI = [0.04, 0.36]), N1 (F_(1,53.87)_ = 47.88, *p* < 0.001, η_p_^2^ = 0.47, 95% CI = [0.28, 0.61]) and P2 (F_(1,54)_ = 154.14, *p* < 0.001, η_p_^2^ = 0.74, 95% CI = [0.62, 0.82]). The age difference for P1 was present in the left hemisphere only (t_(69.8)_ = 4.16, *p* < 0.001, d = 1.58, 95% CI = [0.81, 2.35]), as we reported no age-related modulation of P1 for the right hemisphere (t_(70)_ = 2.36, *p* = 0.10).

When comparing reorientation and control tasks, there was no difference in P1 latency (F_(1,161.15)_ = 0.08, *p* = 0.77). We observed a similar pattern for the other two components, with a later peak for the reorientation task for N1 (F_(1,161.01)_ = 9.34, *p* = 0.003, η_p_^2^ = 0.05, 95% CI = [0.01, 0.14]) and P2 (F_(1,162)_ = 58.59, *p* < 0.001, η_p_^2^ = 0.27, 95% CI = [0.16, 0.37]).

For lateralization, comparing left and right hemispheres, we observed a later P1 peak for the left hemisphere (F_(1,161.15)_ = 4.96, *p* = 0.027, η_p_^2^ = 0.03, 95% CI = [0.00, 0.10]), with a similar pattern for the N1 component (F_(1,161.01)_ = 6.31, *p* = 0.013, η_p_^2^ = 0.04, 95% CI = [0.00, 0.11]). Considering P2, we found no difference between the left and right hemispheres (F_(1,162)_ = 1.17, *p* = 0.28). The later P1 peak was observed for older adults only (t_(161.2)_ = 3.39, *p* = 0.005, d = 0.64, 95% CI = [0.26, 1.02]).

### 3.4 ERSP results comparing reorientation and control tasks

Finally, we examined brain oscillations by computing ERSP. This allowed us to examine additional information about the underlying cognitive process beyond ERPs (Herrmann et al., 2014).

For all ERSP analyses (**Figure 5.A**.), we found a similar pattern consisting of an increased synchronization (*i.e.,* an increase in power interpreted as an increase in neural firing synchronization of the underlying neuronal population compared to the selected baseline) in low frequency bands, *delta*/*theta* (1-8 Hz), occurring 50 ms before image presentation for young adults and lasting up to +400 ms and up to +500 ms for older adults in the reorientation task. This synchronization was followed by desynchronization in the higher frequency bands *alpha* (8-13 Hz) and *beta* (13-30 Hz). We then conducted a cluster-based permutation tests on these results (**Figure 5.B.**). In the right hemisphere, we reported a decreased synchronization in the *alpha/beta* band in older adults compared to young adults, starting from +200 ms and lasting until +1000 ms. In the left hemisphere, this result was significant only within a short window around +200 ms. We also reported a decreased synchronization in the *delta* (1-3 Hz) band for older adults compared to their younger counterparts, starting from 50 ms before image presentation up to +200 ms. Older adults also showed an increase in *theta* (3-8 Hz) synchronization, with a burst starting just after +200 ms and lasting up to +400 ms, and a decrease in high *beta* band synchronization for the reorientation task specific to the right hemisphere.

**Figure 5.**
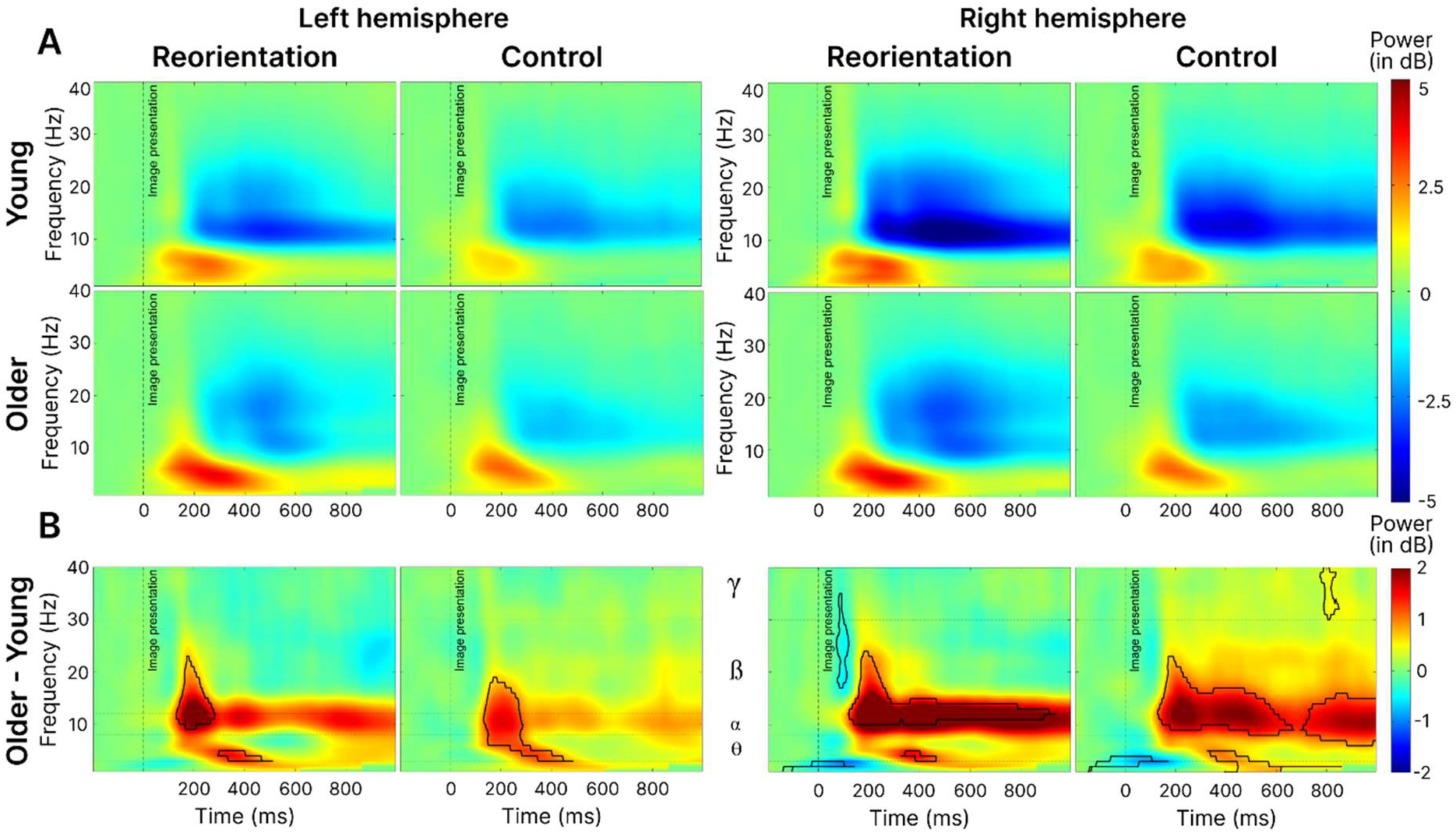
ERSP results comparing reorientation and control tasks. **A.** Grand-average ERSPs, considering age (young/older), task (reorientation/control) and hemisphere (left/right) as variables, using decibel baseline (-150 to -50 ms) normalization. Activity averaged over P6-P8-PO8 electrodes for the right hemisphere and over P5-P7-PO7 electrodes for the left hemisphere. **B.** ERSP activity of older minus young adults. The black line corresponds to cluster-based permutation tests results with p-value < 0.05.

### 3.5 Correlation analyses between ERP and Behavioral data

We performed correlational analyses between ERP (peak amplitudes and latencies) and behavioral data (reaction times and accuracies), separately for young and older adults. This resulted in a total of 48 comparisons, with no significant results after Bonferroni correction. A table with all the uncorrected *p*- values is available in the Supplementary Materials (Table S2).

## 4. Discussion

This EEG study aimed to investigate age-related differences during a landmark-based reorientation task and their neural correlates. Consistent with previous studies, our results indicate that older adults demonstrate reduced navigational abilities, as evidenced by slower and less precise reorientation. However, in contrast with our initial hypothesis, the perception of landmarks of different sizes was not a deteriorating factor for older adults’ performance on the task, even though it was associated with a delayed P1 component in older adults only. Age-related reorientation deficits were associated with differences in the neural dynamics of high-level visual processing. Indeed, we found delayed latencies of the P1, N1 and P2 components recorded from occipito-parietal electrodes associated with scene-selective regions. Moreover, older adults displayed decreased P1 and P2 amplitudes as well as lower *alpha*/*beta* desynchronization in the right hemisphere specifically. Finally, only older adults exhibited increased N1 amplitude in both hemispheres for the reorientation task, accompanied by higher levels of *theta* power. These results points toward a three-part process that may contribute to older adults’ difficulties in landmark reorientation, involving slower and less efficient visual processing, more effortful processing of visuospatial information, and a deficit in the attentional mechanism related to the selection of task-relevant stimuli.

### Reorientation performance is impaired in aging and reaction time decreases similarly for both age groups when perceptual difficulty increases

The behavioral results indicated that young adults performed better, with a lower reaction time and a higher rate of recovered paths. These results support previous findings that older adults are impaired during a navigation task which solely relies on the use of visual landmarks for reorientation (Bécu et al., 2023; West et al., 2023). It is worth mentioning, that despite their diminished performance, older adults still achieved a high level of accuracy which may be explained by the relative simplicity of the task and their healthy cognitive status. Increasing perceptual difficulty (*i.e.,* decreasing landmark size) did not increase error rate, but it did lead to an increase in reaction time in both age groups. This finding is consistent with the well-characterized relationship between stimulus size and reaction time (Plewan & Rinkenauer, 2017; Sperandio et al., 2009). However, contrary to our hypotheses, older adults did not show greater difficulty than young adults when perceptual difficulty was manipulated. This lack of behavioral difference is in line with our EEG findings, which showed no interaction effect between perceptual difficulty and age; both age groups displayed a similar increase in P1 amplitude in the small landmark condition. However, we reported a later P1 peak amplitude only for older adults in the small condition compared to the large condition. It has been suggested that the P1 component is modulated by visuospatial attention (Di Russo et al., 2003), but the exact functional basis of this effect remains debated. The “P1 inhibition-timing” model proposed by Klimesch et al. (2007, 2011) suggests that the P1 reflects a transient inhibitory filter that operates to increase the signal-to-noise ratio allowing an efficient early categorization process. This component may be the earliest index of attentional control, being increased over posterior regions when processing complexity is high (Fellinger et al., 2012). Moreover, the fact that older adults presented a decreased P1 amplitude suggests that they may have impairments in the early categorization process of visual stimuli which are exacerbated by the reorientation task. Then, when faced with increased perceptual difficulty older adults required more time to allocate greater attentional resources to the harder perceptual task (Sawetsuttipan et al., 2023). These findings align with the two leading cognitive aging hypotheses which posit that aging is associated with an inhibitory deficit (*i.e.,* decreased P1 amplitude) and the processing speed hypothesis, (*i.e*., increased P1 latency) when the perceptual difficulty of the task is increased (Finnigan et al., 2011; Gazzaley et al., 2008). It is worth noting that although there was a difference in P1 latency, neither age-related modulation was observed for the later components (namely, N1 and P2) nor for behavioral data. This suggests that the delay in P1 latency may not significantly affect performance with a possible later mechanism compensating for this early delay.

### N1 amplitude and theta power reflects increased resources allocation for landmark processing by older adults

Regarding the N1 component, we observed that older adults exhibited an increased amplitude in both hemispheres compared to young adults. Additionally, we found that older adults had a higher N1 amplitude during the reorientation phase than during passive perception, and that this pattern was not observed in young adults. Some authors have suggested that the N1 component reflect enhanced visual information processing (Luck, 1995; Vogel & Luck, 2000) and visual stimuli discrimination (Finnigan et al., 2011; Hillyard et al., 1998; Hopf et al., 2002; Warbrick et al., 2014; Wiegand et al., 2014). The N1 was previously associated with the activity of the posterior precuneus (Natale et al., 2006). This region plays a role in allocating attention to spatial information, encoding and retrieving spatial memories, and identifying and using relevant landmarks (Cona & Scarpazza, 2019; Delaux et al., 2021). Intracranial EEG recordings also pointed out the activity of the precuneus activity occurring around 210ms after the presentation of a stimulus, while also highlighting the activity of the PPA around 170ms (Bastin et al., 2013; Vlcek et al., 2020). During the same time window as the N1 component, we also observed an increase in *theta* activity. The link between *theta* power and the N1 component was proposed by Klimesch et al. (2004) arguing that the power in the N1 time window was generated primarily by frequencies in the *theta* range (Gruber et al., 2005; Van der Lubbe et al., 2016). Here, we observed an increased *theta* synchronization during the reorientation task, which was more pronounced in older adults, with a burst occurring after 200ms. This increase in *theta* activity was also observed by Lithfous et al. (2018) during a maze reorientation task and associated with better performance in the older group. They interpreted their results as a possible compensatory mechanism exhibited by high-performing older adults, which highlights the importance of parietal *theta* activity for a successful visually-guided navigation (Chrastil et al., 2022). Taken together our results emphasize the role of the N1 component in landmark-based spatial navigation in the context of aging. They provide evidence for an increased bilateral activity in the *theta* band on electrodes associated with scene-selective regions suggesting that older adults resort to more extensive neural resources to process visual landmarks.

### Age-related decrements in selective attention during task-relevant information processing

Regarding the P2 component, we found that older adults exhibited a reduced amplitude compared to younger adults. Conversely, we observed a similar enhancement of amplitude in both age groups when they performed the reorientation task versus the passive perception. These results seem to indicate that the age-related decrease in P2 amplitude reflects a general impairment in the capacity to complete a visual discrimination task, regardless of the reorientation task. The literature has proposed that this posterior P2 component may reflect the mediation of information between memory systems, as a way to compare visual inputs and information stored in the working memory (Cepeda-Freyre et al., 2020; Freunberger et al., 2007; Lefebvre et al., 2005). Other recent findings suggest that the P2 component may rather represent a top-down attentional control during visual processing of objects (Lai et al., 2020), indicating enhanced selective attention by task-relevant stimulus (Freunberger et al., 2007; Mecklinger et al., 2009; Philips & Takeda, 2009). As we observed a similar age-related decrease in a task that did not involve working memory (*i.e*., a passive perception), we argue that the P2 amplitude decrease we report may reflect the well-supported age-related decline in top-down selection of task-relevant objects, which are the landmarks in our case (Lai et al., 2020). However, we cannot exclude that this decrease in P2 amplitude among older adults could also be ascribed to the decline in spatial working memory due to cognitive aging as reported in previous studies (Finnigan et al., 2011; Klencklen et al., 2012). This proposed age-related decrease in spatial information processing is also supported by our ERSP results, showing a decrease of *alpha/beta* desynchronization among older adults. It has been suggested that these frequency bands support the endogenous activation of neuronal ensembles involved in task-relevant information processing (Griffiths et al., 2019; Hanslmayr et al., 2012; Spitzer & Haegens, 2017). They were also previously observed during good reorientation choices in spatial navigation task (Chrastil et al., 2022), interpreted as the reflect of memory retrieval process (Klimesch, 1997, 1999). Finally, we hypothesized an increase in P2 amplitude with aging, in light of the findings from Ramanoël et al. (2020) who reported an increase in OPA activity among older adults during active reorientation, the OPA activity proposed to be reflected in the parietal P2 component (Harel et al., 2016, 2022; Kaiser et al., 2020). However, in their work, Ramanoël et al. (2020) had subjects actively navigate in the maze, and used fMRI recordings, which have some important differences that may account for the differences we observed. Moreover, the OPA may not be the only scene-selective brain regions contributing to the P2 component, and spatial sensitivity of scalp EEG does not allow us to distinguish between OPA and PPA for example. This was suggested by Kaiser et al. (2020) who also reported, along with OPA, activity of the PPA during this time window which was proposed in an iEEG study to last for thousands of milliseconds (Vlcek et al., 2020), possibly overlapping over occipito-parietal electrodes (Persichetti & Dilks, 2019).

### Aging decreases lateralization of visuospatial processing

Finally, we found a distinct lateralization of brain activity in young adults, with greater activation observed in the right hemisphere than in the left hemisphere during both the passive perception and reorientation phases of the experiment. Harel et al. (2016, 2022) similarly observed higher amplitudes in the right hemisphere among young adults only, during a passive scene perception task. This is consistent with the commonly accepted notion that selective spatial attention and spatial working memory are controlled by a predominantly right hemisphere network (Awh & Jonides, 2001; Young, 2012). This also appears to hold true during human spatial navigation as reported in a meta-analysis of 47 fMRI studies (Li et al., 2021). In our results, this lateralization was weaker among older adults for P1 and P2 components, and amplitude was lower in the right hemisphere among older adults only. Using a visuospatial task, Learmonth et al. (2017) also reported decreased right hemisphere control among older adults during a visuospatial task and our results confirm the hypothesis of right hemisphere engagement decrease with age as proposed by the right hemi-aging model (Dolcos et al., 2002). This result also highlights the importance of considering both hemispheres separately when conducting ERP or ERSP investigations of age-related modifications.

### Limitations and perspectives

One of the main limitations of our results is that we did not find any correlation between behavioral and EEG data. This can be explained by the relative behavioral simplicity of our task, which may have prevented us from capturing subtle variations in performance. Furthermore, concerning the modulations of the P1, N1, and P2 components, their interpretation in an independent way could be exaggerated as we acknowledge the possibility that effects observed on later peaks may depend on preceding peaks. For instance, it is conceivable that P1 modulations may exert an effect on N1 peak, rendering the conventional label "component" potentially misleading (Luck, 2005).

In an effort to disentangle reorientation from passive perception, we introduced a cognitive load disparity between the two tasks, which may account in part for the observed results, particularly for the P2 component. To address this issue, future investigations might consider including control tasks relying on stimulus detection and decision-making paradigms, such as the N-back task. Finally, given the changes in visual exploration between young and older adults and their impact on information processing (Bécu et al., 2020, 2023; Durteste et al., 2023; Ryan et al., 2022) it would be worthwhile to investigate the effect of age on gaze patterns during a landmark-based reorientation task, linking EEG with eye-tracking data to gain more insight into how older adults are impaired in using landmarks during reorientation.

## 5. Conclusion

Our study aimed to investigate age-related differences in visuospatial processing and the underlying brain dynamics within scene-selective regions in young and healthy older adults performing a landmark-based reorientation task. Older adults showed reduced reorientation performance along with increased latency of early cortical markers of visual processing in scene-selective regions, suggesting that navigational deficits may result from delayed processing of visuospatial information. Decreasing landmark size and thus increasing perceptual difficulty led to a delayed P1 component only in older adults, suggesting an age-related delayed early categorization of smaller landmarks. Our EEG data also revealed a three-part process that may partially account for older adults’ challenges during landmark reorientation. First, a delayed and reduced P1 component indicated slower and less efficient visual processing, including stimulus discrimination. Second, the increase in N1 amplitude and theta-band activity indicated a greater demand on cognitive resources, leading to more effortful processing of visuospatial information. Third, the reduction in P2 amplitude associated with *alpha-beta* activity suggested a deficiency in the attentional mechanism for selecting task-relevant stimuli. Finally, our findings underscore the importance of considering both hemispheres separately when studying aging, as they highlight an age-related decrease in right hemisphere specific activity. Taken together, these results highlight the interest of using EEG to gain insight into age-related modulations of neural correlates of visuospatial processing during reorientation, while paving the way for further investigations to better characterize the brain dynamics underlying spatial navigation deficits in healthy older adults.

## CRediT author statement

**Naveilhan Clément**: Conceptualization, Methodology, Formal analysis, Investigation, Writing – Original Draft; **Alexandre Delaux**: Conceptualization, Methodology, Writing – Review & Editing; **Marion Durteste:** Conceptualization, Methodology, Writing - Review & Editing; **Jerome Lebrun:** Writing – Review & Editing, Methodology; **Raphaël Zory:** Writing – Review & Editing, Funding acquisition; **Angelo Arleo:** Writing - Review & Editing, Funding acquisition; **Stephen Ramanoël:** Conceptualization, Supervision, Writing – Review & Editing, Funding Acquisition, Project administration.

## Supporting information

Supplemental Informations

## Acknowledgements

The authors gratefully acknowledge Maud Saulay-Carret for her help with acquisitions and also the voluntary participants involved in this research.

This work was supported by the French government through the UCA^JEDI^ project managed by the National Research Agency (ANR-15- IDEX-01) and, in particular, by the interdisciplinary Institute for Modeling in Neuroscience and Cognition (NeuroMod) of Université Côte d’Azur. It was also supported by the Fondation pour la Recherche sur Alzheimer, the French National Research Agency (ANR-18- CHIN-0002), the LabEx LIFESENSES (ANR-10-LABX-65), and the IHU FOReSIGHT (ANR-18-IAHU-01). The funders had no role in study design, data collection and analysis, decision to publish or preparation of the manuscript.

## Code and data availability

All raw data, codes and stimuli generated for the purpose of analyses in this paper are available via the OSF repository: https://osf.io/crne8/?view_only=7a34edf4644e49a796442388ce7ac885

## References

Akaike, H. (1974). A new look at the statistical model identification. IEEE Transactions on Automatic Control, 19(6), 716-723. 10.1109/TAC.1974.1100705

Awh, E., & Jonides, J. (2001). Overlapping mechanisms of attention and spatial working memory. Trends in Cognitive Sciences, 5(3), 119-126. 10.1016/S1364-6613(00)01593-X

Barrash, J. (1994). Age-related decline in route learning ability. Developmental Neuropsychology, 10(3), 189-201. 10.1080/87565649409540578

Bastin, J., Vidal, J. R., Bouvier, S., Perrone-Bertolotti, M., Bénis, D., Kahane, P., David, O., Lachaux, J.-P., & Epstein, R. A. (2013). Temporal Components in the Parahippocampal Place Area Revealed by Human Intracerebral Recordings. Journal of Neuroscience, 33(24), 10123-10131. 10.1523/JNEUROSCI.4646-12.2013

Bates, D., Mächler, M., Bolker, B., & Walker, S. (2014). *Fitting Linear Mixed-Effects Models using lme4* (arXiv:1406.5823; Version 1). arXiv. 10.48550/arXiv.1406.5823

Bécu, M., Sheynikhovich, D., Ramanoël, S., Tatur, G., Ozier-Lafontaine, A., Authié, C. N., Sahel, J.- A., & Arleo, A. (2023). Landmark-based spatial navigation across the human lifespan. eLife, 12, e81318. 10.7554/eLife.81318

Bécu, M., Sheynikhovich, D., Tatur, G., Agathos, C. P., Bologna, L. L., Sahel, J.-A., & Arleo, A. (2020). Age-related preference for geometric spatial cues during real-world navigation. Nature Human Behaviour, 4(1), Article 1. 10.1038/s41562-019-0718-z

Bigdely-Shamlo, N., Mullen, T., Kothe, C., Su, K.-M., & Robbins, K. A. (2015). The PREP pipeline : Standardized preprocessing for large-scale EEG analysis. Frontiers in Neuroinformatics, 9, 16. 10.3389/fninf.2015.00016

Bohbot, V. D., Copara, M. S., Gotman, J., & Ekstrom, A. D. (2017). Low-frequency theta oscillations in the human hippocampus during real-world and virtual navigation. Nature Communications, 8, 14415. 10.1038/ncomms14415

Bonner, M. F., & Epstein, R. A. (2017). Coding of navigational affordances in the human visual system. PNAS Proceedings of the National Academy of Sciences of the United States of America, 114, 4793-4798. 10.1073/pnas.1618228114

Burns, P. C. (1999). Navigation and the mobility of older drivers. The Journals of Gerontology. Series B, Psychological Sciences and Social Sciences, 54(1), S49–55. 10.1093/geronb/54b.1.s49

Cepeda-Freyre, H. A., Garcia-Aguilar, G., Eguibar, J. R., & Cortes, C. (2020). Brain Processing of Complex Geometric Forms in a Visual Memory Task Increases P2 Amplitude. Brain Sciences, 10(2), 114. 10.3390/brainsci10020114

Chang, C.-Y., Hsu, S.-H., Pion-Tonachini, L., & Jung, T.-P. (2020). Evaluation of Artifact Subspace Reconstruction for Automatic Artifact Components Removal in Multi-Channel EEG Recordings. IEEE Transactions on Biomedical Engineering, 67(4), 1114-1121. 10.1109/TBME.2019.2930186

Chrastil, E. R., Rice, C., Goncalves, M., Moore, K. N., Wynn, S. C., Stern, C. E., & Nyhus, E. (2022). Theta oscillations support active exploration in human spatial navigation. NeuroImage, 262, 119581. 10.1016/j.neuroimage.2022.119581

Cohen, M. X. (2014). Analyzing Neural Time Series Data : Theory and Practice. MIT Press.

Cona, G., & Scarpazza, C. (2019). Where is the « where » in the brain? A meta-analysis of neuroimaging studies on spatial cognition. Human Brain Mapping, 40(6), 1867-1886. 10.1002/hbm.24496

Coughlan, G., Laczó, J., Hort, J., Minihane, A.-M., & Hornberger, M. (2018). Spatial navigation deficits—Overlooked cognitive marker for preclinical Alzheimer disease? Nature Reviews. Neurology, 14(8), 496-506. 10.1038/s41582-018-0031-x

de Condappa, O., & Wiener, J. M. (2014). Human place and response learning : Navigation strategy selection, pupil size and gaze behavior. Psychological Research, 80(1), 82-93. 10.1007/s00426-014-0642-9

Delaux, A., de Saint Aubert, J.-B., Ramanoël, S., Bécu, M., Gehrke, L., Klug, M., Chavarriaga, R., Sahel, J.-A., Gramann, K., & Arleo, A. (2021). Mobile brain/body imaging of landmark-based navigation with high-density EEG. European Journal of Neuroscience, 54(12), 8256-8282. 10.1111/ejn.15190

Delorme, A., & Makeig, S. (2004). EEGLAB : An open source toolbox for analysis of single-trial EEG dynamics including independent component analysis. Journal of Neuroscience Methods, 134(1), 9-21. 10.1016/j.jneumeth.2003.10.009

Di Russo, F., Martínez, A., & Hillyard, S. A. (2003). Source Analysis of Event-related Cortical Activity during Visuo-spatial Attention. Cerebral Cortex, 13(5), 486-499. 10.1093/cercor/13.5.486

Dilks, D. D., Julian, J. B., Paunov, A. M., & Kanwisher, N. (2013). The Occipital Place Area Is Causally and Selectively Involved in Scene Perception. Journal of Neuroscience, 33(4), 1331-1336. 10.1523/JNEUROSCI.4081-12.2013

Dolcos, F., Rice, H. J., & Cabeza, R. (2002). Hemispheric asymmetry and aging : Right hemisphere decline or asymmetry reduction. Neuroscience & Biobehavioral Reviews, 26(7), 819-825. 10.1016/S0149-7634(02)00068-4

Durteste, M., Delaux, A., Ariztégui, A., Cottereau, B., Sheynikhovich, D., Ramanoel, S., & Arleo, A. (2023). Age-related disparities in oscillatory dynamics within scene-selective regions during spatial navigation. 10.1101/2023.10.16.562507

Ekstrom, A. D. (2015). Why vision is important to how we navigate. Hippocampus, 25(6), 731-735. 10.1002/hipo.22449

Ekstrom, A. D., Huffman, D. J., & Starrett, M. (2017). Interacting networks of brain regions underlie human spatial navigation : A review and novel synthesis of the literature. Journal of Neurophysiology, 118(6), 3328-3344. 10.1152/jn.00531.2017

Epstein, R. A., Patai, E. Z., Julian, J. B., & Spiers, H. J. (2017). The cognitive map in humans : Spatial navigation and beyond. Nature neuroscience, 20(11), 1504-1513. 10.1038/nn.4656

Epstein, R., & Kanwisher, N. (1998). A cortical representation of the local visual environment. Nature, 392(6676), Article 6676. 10.1038/33402

Faubert, J. (2002). Visual perception and aging. Canadian Journal of Experimental Psychology = Revue Canadienne De Psychologie Experimentale, 56(3), 164-176. 10.1037/h0087394

Fellinger, R., Gruber, W., Zauner, A., Freunberger, R., & Klimesch, W. (2012). Evoked traveling alpha waves predict visual-semantic categorization-speed. NeuroImage, 59(4), 3379-3388. 10.1016/j.neuroimage.2011.11.010

Finnigan, S., O’Connell, R. G., Cummins, T. D. R., Broughton, M., & Robertson, I. H. (2011). ERP measures indicate both attention and working memory encoding decrements in aging. Psychophysiology, 48(5), 601-611. 10.1111/j.1469-8986.2010.01128.x

Fischer, L. F., Mojica Soto-Albors, R., Buck, F., & Harnett, M. T. (2020). Representation of visual landmarks in retrosplenial cortex. eLife, 9, e51458. 10.7554/eLife.51458

Foo, P., Warren, W. H., Duchon, A., & Tarr, M. J. (2005). Do Humans Integrate Routes Into a Cognitive MapMap-Versus Landmark-Based Navigation of Novel Shortcuts. *Journal of Experimental Psychology: Learning*, Memory, and Cognition, 31(2), 195-215. 10.1037/0278-7393.31.2.195

Freunberger, R., Klimesch, W., Doppelmayr, M., & Höller, Y. (2007). Visual P2 component is related to theta phase-locking. Neuroscience Letters, 426(3), 181-186. 10.1016/j.neulet.2007.08.062

Friedman, A., Kohler, B., Gunalp, P., Boone, A. P., & Hegarty, M. (2020). A computerized spatial orientation test. Behavior Research Methods, 52(2), 799-812. 10.3758/s13428-019-01277-3

Gazzaley, A., Clapp, W., Kelley, J., McEvoy, K., Knight, R. T., & D’Esposito, M. (2008). Age-related top-down suppression deficit in the early stages of cortical visual memory processing. Proceedings of the National Academy of Sciences of the United States of America, 105(35), 13122-13126. 10.1073/pnas.0806074105

Glover, G. H. (2011). Overview of Functional Magnetic Resonance Imaging. Neurosurgery clinics of North America, 22(2), 133-139. 10.1016/j.nec.2010.11.001

Gramann, K., Hohlefeld, F. U., Gehrke, L., & Klug, M. (2021). Human cortical dynamics during full-body heading changes. Scientific Reports, 11(1), Article 1. 10.1038/s41598-021-97749-8

Greene, M. R., & Oliva, A. (2009). Recognition of natural scenes from global properties : Seeing the forest without representing the trees. Cognitive Psychology, 58(2), 137-176. 10.1016/j.cogpsych.2008.06.001

Griffiths, B. J., Mayhew, S. D., Mullinger, K. J., Jorge, J., Charest, I., Wimber, M., & Hanslmayr, S. (2019). Alpha/beta power decreases track the fidelity of stimulus-specific information. eLife, 8, e49562. 10.7554/eLife.49562

Gruber, W. R., Klimesch, W., Sauseng, P., & Doppelmayr, M. (2005). Alpha Phase Synchronization Predicts P1 and N1 Latency and Amplitude Size. Cerebral Cortex, 15(4), 371-377. 10.1093/cercor/bhh139

Hamid, S. N., Stankiewicz, B., & Hayhoe, M. (2010). Gaze patterns in navigation : Encoding information in large-scale environments. Journal of Vision, 10(12), 28-28. 10.1167/10.12.28

Hanslmayr, S., Staudigl, T., & Fellner, M.-C. (2012). Oscillatory power decreases and long-term memory : The information via desynchronization hypothesis. Frontiers in Human Neuroscience, 6. https://www.frontiersin.org/articles/10.3389/fnhum.2012.00074

Harel, A., Groen, I. I. A., Kravitz, D. J., Deouell, L. Y., & Baker, C. I. (2016). The temporal dynamics of scene processing : A multi-faceted EEG investigation. eNeuro. 10.1523/ENEURO.0139-16.2016

Harel, A., Nador, J. D., Bonner, M. F., & Epstein, R. A. (2022). Early Electrophysiological Markers of Navigational Affordances in Scenes. Journal of Cognitive Neuroscience, 34(3), 397-410. 10.1162/jocn_a_01810

Harris, M. A., & Wolbers, T. (2012). Ageing effects on path integration and landmark navigation. Hippocampus, 22(8), 1770-1780. 10.1002/hipo.22011

Hartmeyer, S., Grzeschik, R., Wolbers, T., & Wiener, J. M. (2017). The Effects of Attentional Engagement on Route Learning Performance in a Virtual Environment : An Aging Study. Frontiers in Aging Neuroscience, 9. https://www.frontiersin.org/articles/10.3389/fnagi.2017.00235

Hayasaka, S., & Nichols, T. E. (2004). Combining voxel intensity and cluster extent with permutation test framework. NeuroImage, 23(1), 54-63. 10.1016/j.neuroimage.2004.04.035

Herrmann, C. S., Rach, S., Vosskuhl, J., & Strüber, D. (2014). Time-frequency analysis of event-related potentials : A brief tutorial. Brain Topography, 27(4), 438-450. 10.1007/s10548-013-0327-5

Hillyard, S. A., Vogel, E. K., & Luck, S. J. (1998). Sensory gain control (amplification) as a mechanism of selective attention : Electrophysiological and neuroimaging evidence. Philosophical Transactions of the Royal Society of London. Series B, Biological Sciences, 353(1373), 1257-1270. 10.1098/rstb.1998.0281

Hopf, J.-M., Vogel, E., Woodman, G., Heinze, H.-J., & Luck, S. J. (2002). Localizing visual discrimination processes in time and space. Journal of Neurophysiology, 88(4), 2088-2095. 10.1152/jn.2002.88.4.2088

Jacobs, J. (2014). Hippocampal theta oscillations are slower in humans than in rodents : Implications for models of spatial navigation and memory. *Philosophical Transactions of the Royal Society of London. Series B*, Biological Sciences, 369(1635), 20130304. 10.1098/rstb.2013.0304

Jacobs, J., Korolev, I. O., Caplan, J. B., Ekstrom, A. D., Litt, B., Baltuch, G., Fried, I., Schulze-Bonhage, A., Madsen, J. R., & Kahana, M. J. (2010). Right-lateralized Brain Oscillations in Human Spatial Navigation. Journal of Cognitive Neuroscience, 22(5), 824-836. 10.1162/jocn.2009.21240

Janzen, G., & van Turennout, M. (2004). Selective neural representation of objects relevant for navigation. Nature Neuroscience, 7(6), Article 6. 10.1038/nn1257

Julian, J. B., Keinath, A. T., Marchette, S. A., & Epstein, R. A. (2018). The Neurocognitive Basis of Spatial Reorientation. Current Biology, 28(17), R1059-R1073. 10.1016/j.cub.2018.04.057

Kaiser, D., Häberle, G., & Cichy, R. M. (2020). Cortical sensitivity to natural scene structure. Human Brain Mapping, 41(5), 1286-1295. 10.1002/hbm.24875

Kalafat, M., Hugonot-Diener, L., & Poitrenaud, J. (2003). Standardisation et étalonnage français du « Mini Mental State » (MMS) version GRÉCO. [French standardization and range for the GRECO version of the « Mini Mental State » (MMS).]. Revue de Neuropsychologie, 13, 209-236.

Klencklen, G., Després, O., & Dufour, A. (2012). What do we know about aging and spatial cognition? Reviews and perspectives. Ageing Research Reviews, 11(1), 123-135. 10.1016/j.arr.2011.10.001

Klimesch, W. (1997). EEG-alpha rhythms and memory processes. International Journal of Psychophysiology: Official Journal of the International Organization of Psychophysiology, 26(1-3), 319-340. 10.1016/s0167-8760(97)00773-3

Klimesch, W. (1999). EEG alpha and theta oscillations reflect cognitive and memory performance : A review and analysis. Brain Research Reviews, 29(2), 169-195. 10.1016/S0165-0173(98)00056-3

Klimesch, W. (2011). Evoked alpha and early access to the knowledge system : The P1 inhibition timing hypothesis. Brain Research, 1408, 52-71. 10.1016/j.brainres.2011.06.003

Klimesch, W., Sauseng, P., & Hanslmayr, S. (2007). EEG alpha oscillations : The inhibition–timing hypothesis. Brain Research Reviews, 53(1), 63-88. 10.1016/j.brainresrev.2006.06.003

Klimesch, W., Schack, B., Schabus, M., Doppelmayr, M., Gruber, W., & Sauseng, P. (2004). Phase-locked alpha and theta oscillations generate the P1–N1 complex and are related to memory performance. Cognitive Brain Research, 19(3), 302-316. 10.1016/j.cogbrainres.2003.11.016

Klug, M., & Gramann, K. (2021). Identifying key factors for improving ICA-based decomposition of EEG data in mobile and stationary experiments. European Journal of Neuroscience, 54(12), 8406-8420. 10.1111/ejn.14992

Klug, M., & Kloosterman, N. A. (2022). Zapline-plus : A Zapline extension for automatic and adaptive removal of frequency-specific noise artifacts in M/EEG. Human Brain Mapping, 43(9), 2743-2758. 10.1002/hbm.25832

Koch, C., Li, S. C., Polk, T. A., & Schuck, N. W. (2020). Effects of aging on encoding of walking direction in the human brain. Neuropsychologia. 10.1016/j.neuropsychologia.2020.107379

Kothe, C. A. E., & Jung, T.-P. (2015). Artifact removal techniques with signal reconstruction (World Intellectual Property Organization Brevet WO2015047462A9). https://patents.google.com/patent/WO2015047462A9/en

Kravitz, D. J., Peng, C. S., & Baker, C. I. (2011). Real-World Scene Representations in High-Level Visual Cortex : It’s the Spaces More Than the Places. Journal of Neuroscience, 31(20), 7322-7333. 10.1523/JNEUROSCI.4588-10.2011

Kropotov, J., Ponomarev, V., Tereshchenko, E. P., Müller, A., & Jäncke, L. (2016). Effect of Aging on ERP Components of Cognitive Control. Frontiers in Aging Neuroscience, 8, 69. 10.3389/fnagi.2016.00069

Lai, L. Y., Frömer, R., Festa, E. K., & Heindel, W. C. (2020). Age-related changes in the neural dynamics of bottom-up and top-down processing during visual object recognition : An electrophysiological investigation. Neurobiology of Aging, 94, 38-49. 10.1016/j.neurobiolaging.2020.05.010

Learmonth, G., Benwell, C. S. Y., Thut, G., & Harvey, M. (2017). Age-related reduction of hemispheric lateralisation for spatial attention : An EEG study. NeuroImage, 153, 139-151. 10.1016/j.neuroimage.2017.03.050

Lefebvre, C. D., Marchand, Y., Eskes, G. A., & Connolly, J. F. (2005). Assessment of working memory abilities using an event-related brain potential (ERP)-compatible digit span backward task. Clinical Neurophysiology: Official Journal of the International Federation of Clinical Neurophysiology, 116(7), 1665-1680. 10.1016/j.clinph.2005.03.015

Lester, A. W., Moffat, S. D., Wiener, J. M., Barnes, C. A., & Wolbers, T. (2017). The Aging Navigational System. Neuron, 95(5), 1019-1035. 10.1016/j.neuron.2017.06.037

Li, J., Zhang, R., Liu, S., Liang, Q., Zheng, S., He, X., & Huang, R. (2021). Human spatial navigation : Neural representations of spatial scales and reference frames obtained from an ALE meta-analysis. NeuroImage, 238, 118264. 10.1016/j.neuroimage.2021.118264

Lithfous, S., Dufour, A., Blanc, F., & Després, O. (2014). Allocentric but not egocentric orientation is impaired during normal aging : An ERP study. Neuropsychology, 28(5), 761-771. 10.1037/neu0000084

Lithfous, S., Dufour, A., Bouix, C., Pebayle, T., & Després, O. (2018). Reduced parahippocampal theta activity during spatial navigation in low, but not in high elderly performers. Neuropsychology, 32(1), 40-53. 10.1037/neu0000392

Luck, S. (2005). Ten simple rules for designing ERP Experiments. In Event-Related Potentials : A Methods Handbook (p. 17-32).

Luck, S. J. (1995). Multiple mechanisms of visual-spatial attention : Recent evidence from human electrophysiology. Behavioural Brain Research, *71*(1-2), 113-123. 10.1016/0166-4328(95)00041-0

Luck, S. J. (2014). An Introduction to the Event-Related Potential Technique, second edition. MIT Press.

Maguire, E. (2001). The retrosplenial contribution to human navigation : A review of lesion and neuroimaging findings. Scandinavian Journal of Psychology, 42(3), 225-238. 10.1111/1467-9450.00233

Makeig, S. (1993). Auditory event-related dynamics of the EEG spectrum and effects of exposure to tones. Electroencephalography and Clinical Neurophysiology, 86(4), 283-293. 10.1016/0013-4694(93)90110-h

Marchette, S. A., Vass, L. K., Ryan, J., & Epstein, R. A. (2014). Anchoring the neural compass : Coding of local spatial reference frames in human medial parietal lobe. Nature Neuroscience, 17(11), Article 11. 10.1038/nn.3834

Maris, E., & Oostenveld, R. (2007). Nonparametric statistical testing of EEG-and MEG-data. Journal of Neuroscience Methods, 164(1), 177-190. 10.1016/j.jneumeth.2007.03.024

Mecklinger, A., Parra, M., & Waldhauser, G. T. (2009). ERP correlates of intentional forgetting. Brain Research, 1255, 132-147. 10.1016/j.brainres.2008.11.073

Moca, V. V., Bârzan, H., Nagy-Dăbâcan, A., & Mureșan, R. C. (2021). Time-frequency super-resolution with superlets. Nature Communications, 12(1), Article 1. 10.1038/s41467-020-20539-9

Mueller, V., Brehmer, Y., von Oertzen, T., Li, S.-C., & Lindenberger, U. (2008). Electrophysiological correlates of selective attention : A lifespan comparison. BMC Neuroscience, 9(1), 18. 10.1186/1471-2202-9-18

Natale, E., Marzi, C. A., Girelli, M., Pavone, E. F., & Pollmann, S. (2006). ERP and fMRI correlates of endogenous and exogenous focusing of visual-spatial attention. The European Journal of Neuroscience, 23(9), 2511-2521. 10.1111/j.1460-9568.2006.04756.x

Oostenveld, R., Fries, P., Maris, E., & Schoffelen, J.-M. (2011). FieldTrip : Open source software for advanced analysis of MEG, EEG, and invasive electrophysiological data. Computational Intelligence and Neuroscience, 2011, 156869. 10.1155/2011/156869

Oostenveld, R., & Oostendorp, T. F. (2002). Validating the boundary element method for forward and inverse EEG computations in the presence of a hole in the skull. Human Brain Mapping, 17(3), 179-192. 10.1002/hbm.10061

Palmer, J. A., Makeig, S., Kreutz-Delgado, K., & Rao, B. D. (2008). Newton method for the ICA mixture model. 2008 IEEE International Conference on Acoustics, Speech and Signal Processing, 1805-1808. 10.1109/ICASSP.2008.4517982

Persichetti, A. S., & Dilks, D. D. (2018). Dissociable Neural Systems for Recognizing Places and Navigating through Them. Journal of Neuroscience, 38(48), 10295-10304. 10.1523/JNEUROSCI.1200-18.2018

Persichetti, A. S., & Dilks, D. D. (2019). Distinct representations of spatial and categorical relationships across human scene-selective cortex. Proceedings of the National Academy of Sciences, 116(42), 21312-21317. 10.1073/pnas.1903057116

Philips, S., & Takeda, Y. (2009). An EEG/ERP study of efficient versus inefficient visual search. Proceedings of the Annual Meeting of the Cognitive Science Society, 31(31). https://escholarship.org/uc/item/93x9f8bb

Pion-Tonachini, L., Kreutz-Delgado, K., & Makeig, S. (2019). ICLabel : An automated electroencephalographic independent component classifier, dataset, and website. NeuroImage, 198, 181-197. 10.1016/j.neuroimage.2019.05.026

Plewan, T., & Rinkenauer, G. (2017). Simple reaction time and size–distance integration in virtual 3D space. Psychological Research, 81(3), 653-663. 10.1007/s00426-016-0769-y

Ramanoël, S., Durteste, M., Bécu, M., Habas, C., & Arleo, A. (2020). Differential Brain Activity in Regions Linked to Visuospatial Processing During Landmark-Based Navigation in Young and Healthy Older Adults. Frontiers in Human Neuroscience, 14. https://www.frontiersin.org/articles/10.3389/fnhum.2020.552111

Ramanoël, S., Durteste, M., Bizeul, A., Ozier-Lafontaine, A., Bécu, M., Sahel, J.-A., Habas, C., & Arleo, A. (2022). Selective neural coding of object, feature, and geometry spatial cues in humans. Human Brain Mapping, 43(17), 5281-5295. 10.1002/hbm.26002

Ramanoël, S., Kauffmann, L., Cousin, E., Dojat, M., & Peyrin, C. (2015). Age-related differences in spatial frequency processing during scene categorization. PLoS ONE. 10.1371/journal.pone.0134554

Ramanoël, S., York, E., Le Petit, M., Lagrené, K., Habas, C., & Arleo, A. (2019). Age-Related Differences in Functional and Structural Connectivity in the Spatial Navigation Brain Network. Frontiers in Neural Circuits, 13. https://www.frontiersin.org/articles/10.3389/fncir.2019.00069

Ryan, J. D., Wynn, J. S., Shen, K., & Liu, Z.-X. (2022). Aging changes the interactions between the oculomotor and memory systems. *Neuropsychology, Development, and Cognition. Section B, Aging*, Neuropsychology and Cognition, 29(3), 418-442. 10.1080/13825585.2021.2007841

Sawetsuttipan, P., Phunchongharn, P., Ounjai, K., Salazar, A., Pongsuwan, S., Intrachooto, S., Serences, J. T., & Itthipuripat, S. (2023). Perceptual Difficulty Regulates Attentional Gain Modulations in Human Visual Cortex. The Journal of Neuroscience: The Official Journal of the Society for Neuroscience, 43(18), 3312-3330. 10.1523/JNEUROSCI.0519-22.2023

Segen, V., Avraamides, M. N., Slattery, T. J., & Wiener, J. M. (2021). Age-related differences in visual encoding and response strategies contribute to spatial memory deficits. Memory & Cognition, 49(2), 249-264. 10.3758/s13421-020-01089-3

Silson, E. H., Steel, A. D., & Baker, C. I. (2016). Scene-Selectivity and Retinotopy in Medial Parietal Cortex. Frontiers in Human Neuroscience, 10, 412. 10.3389/fnhum.2016.00412

Sperandio, I., Savazzi, S., Gregory, R. L., & Marzi, C. A. (2009). Visual Reaction Time and Size Constancy. Perception, 38(11), 1601-1609. 10.1068/p6421

Spitzer, B., & Haegens, S. (2017). Beyond the Status Quo : A Role for Beta Oscillations in Endogenous Content (Re)Activation. eNeuro, 4(4). 10.1523/ENEURO.0170-17.2017

Srokova, S., Hill, P. F., Koen, J. D., King, D. R., & Rugg, M. D. (2020). Neural differentiation is moderated by age in scene-selective, but not face-selective, cortical regions. eNeuro, 7(3). 10.1523/ENEURO.0142-20.2020

Sun, L., Frank, S. M., Epstein, R. A., & Tse, P. U. (2021). The parahippocampal place area and hippocampus encode the spatial significance of landmark objects. NeuroImage, 236, 118081. 10.1016/j.neuroimage.2021.118081

Van der Lubbe, R. H. J., Szumska, I., & Fajkowska, M. (2016). Two Sides of the Same Coin : ERP and Wavelet Analyses of Visual Potentials Evoked and Induced by Task-Relevant Faces. Advances in Cognitive Psychology, 12(4), 154-168. 10.5709/acp-0195-3

Vann, S. D., Aggleton, J. P., & Maguire, E. A. (2009). What does the retrosplenial cortex do? Nature Reviews Neuroscience, 10(11), Article 11. 10.1038/nrn2733

Vlcek, K., Fajnerova, I., Nekovarova, T., Hejtmanek, L., Janca, R., Jezdik, P., Kalina, A., Tomasek, M., Krsek, P., Hammer, J., & Marusic, P. (2020). Mapping the Scene and Object Processing Networks by Intracranial EEG. Frontiers in Human Neuroscience, 14, 561399. 10.3389/fnhum.2020.561399

Vogel, E. K., & Luck, S. J. (2000). The visual N1 component as an index of a discrimination process. Psychophysiology, 37(2), 190-203.

Warbrick, T., Arrubla, J., Boers, F., Neuner, I., & Shah, N. J. (2014). Attention to Detail : Why Considering Task Demands Is Essential for Single-Trial Analysis of BOLD Correlates of the Visual P1 and N1. Journal of Cognitive Neuroscience, 26(3), 529-542. 10.1162/jocn_a_00490

Wenczel, F., Hepperle, L., & von Stülpnagel, R. (2017). Gaze behavior during incidental and intentional navigation in an outdoor environment. Spatial Cognition and Computation, *17*(1-2). 10.1080/13875868.2016.1226838

West, G. L., Patai, Z. E., Coutrot, A., Hornberger, M., Bohbot, V. D., & Spiers, H. J. (2023). Landmark-dependent Navigation Strategy Declines across the Human Life-Span : Evidence from Over 37,000 Participants. Journal of Cognitive Neuroscience, 35(3), 452-467. 10.1162/jocn_a_01956

Wiegand, I., Töllner, T., Dyrholm, M., Müller, H. J., Bundesen, C., & Finke, K. (2014). Neural correlates of age-related decline and compensation in visual attention capacity. Neurobiology of Aging, 35(9), 2161-2173. 10.1016/j.neurobiolaging.2014.02.023

Wiener, J., Kmecova, H., & de Condappa, O. (2012). Route repetition and route retracing : Effects of cognitive aging. Frontiers in Aging Neuroscience, 4. https://www.frontiersin.org/articles/10.3389/fnagi.2012.00007

Wolbers, T., & Hegarty, M. (2010). What determines our navigational abilities? Trends in Cognitive Sciences, 14(3), 138-146. 10.1016/j.tics.2010.01.001

Young, A. (2012). Functions of the Right Cerebral Hemisphere. Elsevier.

Zhang, H., Copara, M., & Ekstrom, A. D. (2012). Differential Recruitment of Brain Networks following Route and Cartographic Map Learning of Spatial Environments. PLOS ONE, 7(9), e44886. 10.1371/journal.pone.0044886

