## Supplemental Informations for "Age-related differences in electrophysiological correlates of visuospatial reorientation"

### Supplementary Methods

**Table S1.** Demographic information for the participants, including gender (M = Male, F = Female), and their performance on cognitive evaluations, specifically the Mini Mental State Examination (MMSE) and Spatial Orientation Test (SOT). To assess the differences between age groups for each measure, a Mann-Whitney U-test was employed.

| | **Mean (± SEM)** | | | | |
|---|---|---|---|---|---|
| **SEX (M/F)** | **Young 14/14** | **Older 10/18** | ***p*-value** | **Effect Size** | **CI 95%** |
| **AGE (YEARS OLD)** | 23.93 (± 0.64) | 71.25 (± 1.01) | < 0.001 | $r_{rb}$ = 1.00 | [7.68 ; 13.29] |
| **EDUCATION LEVEL** | 4.11 (± 0.29) | 2.57 (± 0,46) | 0.007 | $r_{rb}$ = 0.41 | [0.19 ; 1.31] |
| **MMSE** | 29.18 (± 0,17) | 29.11 (± 0,21) | 0.972 | $r_{rb}$ = 0.01 | [- 0.52 ; 0.52] |
| **SOT** | 32.63 (± 4.67) | 58.28 (± 5.69) | < 0.001 | $r_{rb}$ = 0.54 | [0.35 ; 1.50] |

As reported in the table above, a notable difference in the level of education between the young and older adult groups was observed, which may be explained by the recruitment of young participants from the local university. This difference is noteworthy, as outlined by Archer et al. (2018), who suggested that the educational level may partially account for the observed neural differences in aging studies. Regarding the spatial orientation test, our results are consistent with those reported by Friedman et al. (2020) for young adults, with mean angular errors ranging between 26 and 37 depending on their experimental conditions. However, concerning the results for the older adults, no other studies reported results with the computerized version for this population. Using the original pencil and paper task, for young adults, showing mean angular errors ranging between 26 and 37, depending on their experimental conditions. However, concerning the results for the older adults, no other studies have reported results with the computerized version for this population. Using the original pencil and paper task, Zancada-Menendez et al. (2016) also reported reduced performance for older adults, but they employed metrics that do not allow a direct comparison with our results. We did not observe any correlations between the EEG or behavioral results and the spatial orientation test results.

**Table S2.** Correlation analysis results, with uncorrected p-values and correlation coefficients for the 48 correlations performed between the ERP and the Behavioral data.

| | ERP measure | Behavior | Uncorrected p-value | Correlation coefficient | 95% Confidence Interval |
|---|---|---|---|---|---|
| Young Right Hemisphere | P1 Latency | Reaction Time | 0.017 | 0.455 | [0.09 ; 0.712] |
| | N1 Latency | | 0.11 | | |
| | P2 Latency | | 0.59 | | |
| | P1 Amplitude | | 0.317 | | |
| | N1 Amplitude | | 0.697 | | |
| | P2 Amplitude | | 0.733 | | |
| | P1 Latency | Accuracy | 0.003 | 0.547 | [0.211 ; 0.767] |
| | N1 Latency | | 0.19 | -0.449 | [-0.709 ; -0.083] |
| | P2 Latency | | 0.732 | | |
| | P1 Amplitude | | 0.69 | | |
| | N1 Amplitude | | 0.61 | | |
| | P2 Amplitude | | 0.756 | | |
| Young Left Hemisphere | P1 Latency | Reaction Time | 0.251 | | |
| | N1 Latency | | 0.019 | -0.442 | [-0.7 ; -0.083] |
| | P2 Latency | | 0.031 | -0.408 | [-0.678 ; 0.041] |
| | P1 Amplitude | | 0.047 | -0.378 | [-0.658 ; -0.006] |
| | N1 Amplitude | | 0.956 | | |
| | P2 Amplitude | | 0.267 | | |
| | P1 Latency | Accuracy | 0.222 | | |
| | N1 Latency | | 0.877 | | |
| | P2 Latency | | 0.762 | | |
| | P1 Amplitude | | 0.31 | | |
| | N1 Amplitude | | 0.675 | | |
| | P2 Amplitude | | 0.412 | | |
| Older Right Hemisphere | P1 Latency | Reaction Time | 0.736 | | |
| | N1 Latency | | 0.241 | | |
| | P2 Latency | | 0.343 | | |
| | P1 Amplitude | | 0.033 | -0.403 | [-0.67 ; -0.035] |
| | N1 Amplitude | | 0.601 | | |
| | P2 Amplitude | | 0.412 | | |
| | P1 Latency | Accuracy | 0.912 | | |
| | N1 Latency | | 0.925 | | |
| | P2 Latency | | 0.463 | | |
| | P1 Amplitude | | 0.326 | | |
| | N1 Amplitude | | 0.954 | | |
| | P2 Amplitude | | 0.588 | | |
| Older Left Hemisphere | P1 Latency | Reaction Time | 0.555 | | |
| | N1 Latency | | 0.753 | | |
| | P2 Latency | | 0.438 | | |
| | P1 Amplitude | | 0.0081 | -0.49 | [-0.730 ; -0.144] |
| | N1 Amplitude | | 0.575 | | |
| | P2 Amplitude | | 0.113 | | |
| | P1 Latency | Accuracy | 0.502 | | |
| | N1 Latency | | 0.04 | 0.39 | [0.019 ; 0.666] |
| | P2 Latency | | 0.421 | | |
| | P1 Amplitude | | 0.36 | | |
| | N1 Amplitude | | 0.845 | | |
| | P2 Amplitude | | 0.941 | | |

#### Supplementary References


Archer, J. A., Lee, A., Qiu, A., & Chen, S.-H. A. (2018). Working memory, age and education : A lifespan fMRI study. *PloS One*, *13*(3), e0194878. https://doi.org/10.1371/journal.pone.0194878

Friedman, A., Kohler, B., Gunalp, P., Boone, A. P., & Hegarty, M. (2020). A computerized spatial orientation test. *Behavior Research Methods*, *52*(2), 799-812. https://doi.org/10.3758/s13428-019-01277-3

Zancada-Menendez, C., Sampedro-Piquero, P., Lopez, L., & McNamara, T. P. (2016). Age and gender differences in spatial perspective taking. *Aging Clinical and Experimental Research*, *28*(2), 289-296. https://doi.org/10.1007/s40520-015-0399-z
